# Nucleoporin Nup153 docks the splicing machinery to the nuclear pore for efficient mRNA processing

**DOI:** 10.1101/2024.09.30.615666

**Authors:** IJ de Castro, L Schuster, C Patiño-Gomez, D Glavas, A Udupa, M Ríos Vázquez, T Symens, G Tulcan, J Heinze, Heras J de las, Robert Reinhardt, Jorge Trojanowski, H Harz, G Stumberger, H Leonhardt, E Schirmer, S Saka, V Laketa, M Lusic

## Abstract

The nuclear pore complex (NPC), composed of proteins termed nucleoporins (Nups), intercalates the nuclear envelope, and is primarily involved in protein trafficking and mRNA export. At the nuclear basket, Nups have been associated with chromatin organization and postulated to function as transcriptional hubs, working in tandem with mRNA export machinery. However, little is known about the intermediate process of RNA splicing at the NPC. Here, we used BioID to screen for interactors of basket-Nups Nup153 and TPR and discovered the enrichment of splicing proteins across all spliceosome complexes (E, A, B, B*, P). The peripheral nature of the interaction between Nup153 and selected splicing components was confirmed by *in-situ* proximity ligation assay and STED microscopy. The presence of splicing components at the NPC, reduced upon splicing inhibition, is partly dependent on Nup153 and functionally correlated to the splicing of Nup153-bound genes. Assessed by DamID, Nup153-bound genes (∼500) are characterized by multiple long introns with lower-than-average GC content. Positioned at the periphery but distinct from the neighbouring lamina-associated domain (LADs) in chromatin signatures and expression levels, these genes showed Nup153-dependent splicing defect, suggesting that splicing occurs at the NPC.

Altogether, our data substantiates the gene gating theory bringing transcription and export, now accompanied by speckle-distant splicing events, at the level of the NPC.

## Introduction

The nuclear space is demarcated physically by the nuclear envelope, composed of the two nuclear membranes and lamina filaments, and intercalated by the nuclear pore complex (NPC). The NPC comprises around 30 proteins, called nucleoporins (Nups), whose canonical function is the trafficking of molecules across the nuclear envelope (*1*). Nups are structurally arranged into a cylindrical scaffold, creating three connected rings: the cytoplasmic and nucleoplasmic ring complexes and the inner ring between them. The large central channel is lined with intrinsically disordered phenylalanine-glycine (FG) rich Nups, that interact with cargo complexes forming a selective barrier and enabling nuclear transport (*2*). At the nuclear side of the NPC, the nuclear basket (NB) protrudes from the nucleoplasmic ring into the nucleoplasm (*3*) with several Nups including Nup98, ELYS, Nup153, and TPR mediating interactions with chromatin. Thus, the NPC represents a multifunctional platform with versatile non-canonical functions (*1*, *4–6*).

In the vicinity of the pore, the NB scaffolding protein TPR maintains the heterochromatin-free space beneath the NPC (*7*), demarcating it from the adjacent lamina environment that harbors large chromatin domains in repressed states (LADs, Lamina-associated domains) (*8*). Unlike repressive LADs the functional relationship between NPC and genome regulation seems to be more varied and less well understood. Most Nups are associated with activated genomic states with few exceptions (*9*, *10*). Nup153, in concert with transcriptional factor Sox2, binds the transcriptional start and termination sites of genes in neuronal cells (*11*). It also orchestrates the binding of Super enhancers (SE) involved in cell type specific regulatory programs (*12*, *13*), organized in phase separated condensates (*13*, *14*). Moreover, the binding of Nup153 to chromatin organizer and insulator CTCF, contributes to genomic organization and transcriptional regulation of early immediate genes at the NPC in response to stimuli (*15*). The NPC thus arises as an important compartment for regulation of certain genes, and concomitantly, depletion of Nup153 in a short time frame by auxin induced degron affects only a few mRNAs (*16*), implying that only a very small fraction of the genome is bound to the NPC at the given time.

Apart from DNA binding functions, machinery assembling at the NPC has been shown to support its proposed role in gene expression during cellular or gene induction, thus contributing to the gene gating theory postulated in 1985 whereby movement of active genes towards the NPC would be favorable for their transcription and export (*17*). In yeast, the NB formation depends on mRNA transcription and subsequent mRNA-protein complex (mRNP) processing (*18*). In human cells, the <u>Tr</u>anscription-<u>Ex</u>port (TREX) complex 2, which couples mRNA transcription and splicing with export (*19*) , associates with the NPC through TPR (*16*, *20*), presumably to facilitate mRNA processing and export following transcription at the NPC (*16*). Although compiling evidence of machinery pertaining to transcription and export has been associated with the NPC, the intermediate process, splicing, has yet to be placed at the level of the NPC.

mRNA splicing happens subsequent to transcription and prior to export, and thus functionally links the two processes occurring mostly co-transcriptionally (*21*). From the early complex E to the A, B, B*, C and post-spliceosome complexes, a total of 200-300 proteins (*22*) participate in spliceosome assembly (*23*). Enrichment of splicing factors is particularly prominent in splicing speckles, that are interchromatin granules in the nucleoplasm of mammalian cells (*24*, *25*). Depending on transcription rates, gene length and number of splice junctions per gene, splicing can occur in proximity to speckles or at different, speckle-distant locations (*26*, *27*) in line with the observed interconnected network of mRNPs distributed throughout the nucleus (*28*, *29*). However, despite the significance of this machinery in mRNA production, splicing interactions with Nups have been far from characterized. Here, to better understand the proposed involvement of the NPC in transcription, we screened for NPC basket interactors by BioID in two cell lines and found enrichment of splicing proteins, with members of all splicing complexes located at the level of the NPC. We correlated the presence of splicing proteins with Nup153 bound genes, mapped here by DamID, and characterized as a subset of cellular genes (∼500) with longer than average introns and lower than average GC content. We postulate that these genes, although distant from splicing speckles, are spliced at the level of the NPC, thus representing a speckle-distant group of spliced genes (*27*).

Altogether, we propose that splicing machinery necessary for effective gene expression programs is found at the NPCs, which together with known assembly of transcription and export machineries at these sites would support the gene gating theory.

## Results

### Splicing machinery is enriched at the Nuclear Basket

We began exploring NB associations with bonafide interactors by targeting Nup153 and TPR using the BioID system in the setting of two cell lines, the commonly used human embryonic kidney fibroblast HEK293T and T lymphocyte Jurkat cells. We screened the interactome of basket Nups by using BirA fused constructs, Nup153-BirA (in HEK293T and Jurkat cells) and TPR-BiraA (in Jurkat cells) and compared them with untagged GFP (in HEK293T& Jurkat cells) (*30*), or with the fusion BirA with LckN18 (*31*) that served as negative or positive controls (HEK293T), respectively (**Fig.S1a**) .

Our stringency criteria for bonafide Nups selected only interactors present in both cell lines, namely triplicates for Nup153 and TPR datasets from HEK293T cells and duplicates for TPR in Jurkats. This resulted in total of 638 confidently detected proteins, henceforth used in subsequent analysis (**Fig. 1a**). Amongst these interactors, we find transport proteins (IPO5/7, TNPO1), transcription machinery (Med1, TADA3, TAF9b, CDK9, NELF, Brd4) or nucleoporins (e.g. Nup35), expected NPC related interactors. STRING analysis on the 638 proteins using 5 clusters (KMeans) and high-confidence interaction scores depicts Splicing and Gene Expression as the two highest scores with False Discovery Rate FDR 1.2^-40^ and FDR 1.2^-25^, respectively (**Fig.1b** and **Fig.S1b**). The top three significantly enriched gene ontology pathways are all related to spliceosome activities, “mRNA splicing”, “spliceosome complex” and “catalytic step 2 spliceosome”, with Benjamini Hochberg corrected p-values to avoid false positives of 5^-16^, 4^-16^ and 4^-8^ (**Fig.1b and 1c)**. In our BioID dataset, members of all spliceosomal complexes were detected, from the early E, to A, B, B* active, C and the post-spliceosome complex P, suggesting a concerted enrichment of the entire splicing machinery at the level of the NPC (**Fig.S1c** and **Fig.1d**) contrasting an exclusive association with late splicing assembly factors loaded concomitantly with the export machinery.

**Figure 1.**
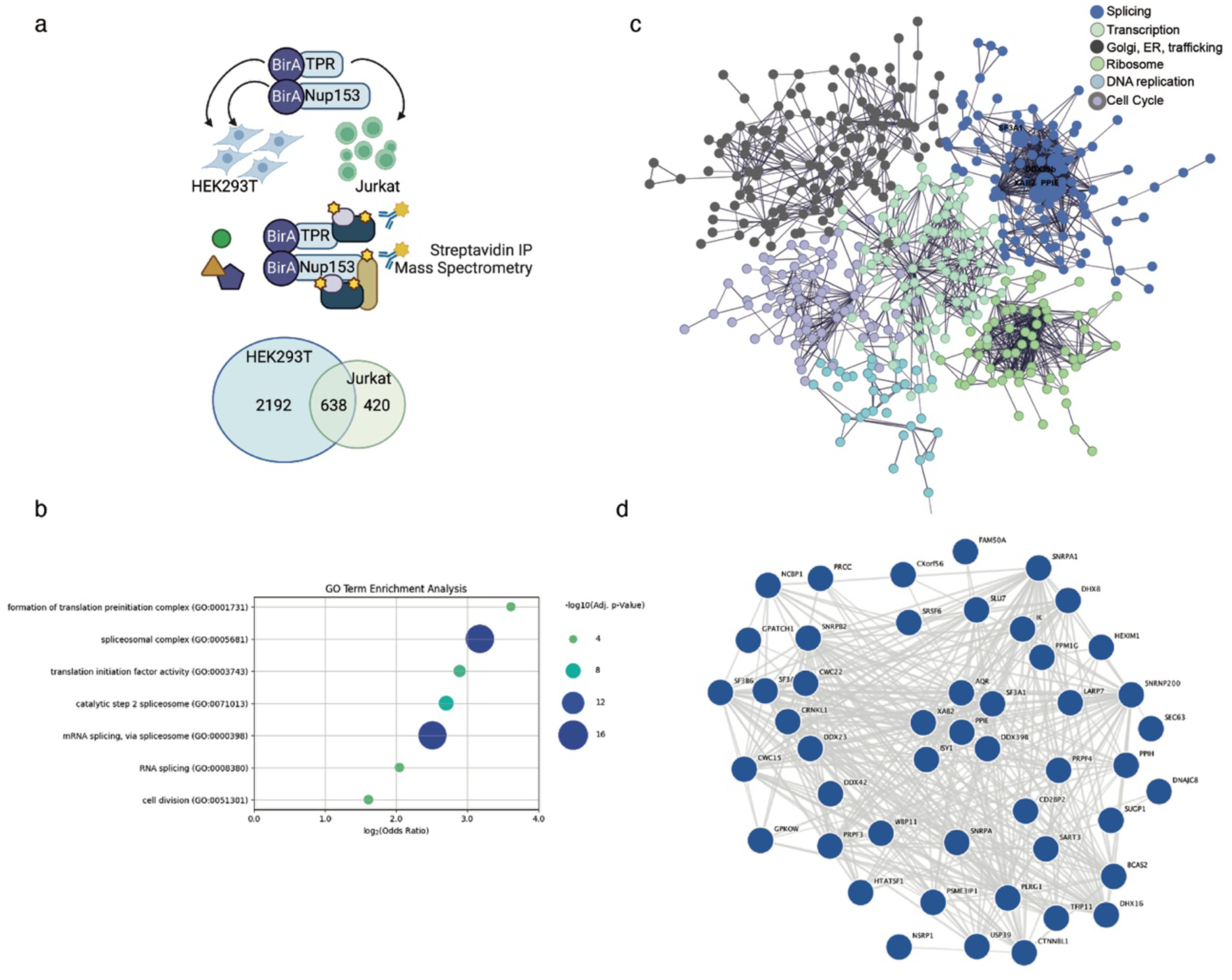
BioID followed by mass spectrometry for nucleoporins TPR and Nup153 reveals the association of splicing proteins at the level of the nuclear basket. **a**. Scheme of the BioID experiment performed in HEK293T and in Jurkat T cells using constructs expressing BirA:Nup153 and/or BirA:TPR. Biotinylated tagged proteins were extracted using streptavidin beads and analysed by Mass-Spectrometry. 638 candidate proteins were narrowed down present in both HEK293T and Jurkat T cell lines datasets. **b**. Gene ontology enrichment analysis indicating the 7 most enriched terms with adjusted p-value and log2 (Odds Ratio). **c**. The 638 proteins were visualised using STRING network analysis, KMeans clustering (5) and high-confidence interaction scores. Clustering indicates proteins enriched across five main gene ontology categories: splicing (dark blue); transcription (light green); Golgi, Endoplasmatic Reticulum and trafficking (grey); ribosome (green), DNA replication (blue) and cell cycle (purple). d. STRING analysis of components of the spliceosome depicted in c. STRING network was exported to Cytoskape and adjusted for visualisation purposes, centred at spliceosome complex members analysed in this study (AQR. XAB2, PPIE, ISY1, SF3A1 and DDX39b).

### Splicing proteins pool at the periphery is in contact with the NPC

In order to investigate further the presence of splicing proteins at the NPC compartment, we have selected members of the Intron Binding Complex (IBC, part of the B* or B-Act complex), the core spliceosome component SF3A1, as well as DDX39b, assembling at the early E complex. We transfected HEK293T cells with eGFP:Nup153 or eGFP and pulled-down the eGFP tag. We observed that four out of five members of the IBC (**Fig.2a** and **Fig.S2a**) are immunoprecipitated with eGFP:Nup153, including AQR, XAB2, PPIE and ISY1. Moreover, Nup153 also immunoprecipitated the early splicing protein DDX39b and the core spliceosome component, SF3A1 (**Fig.2a**). Recently, analysis of splicing machinery components in the 3D nuclear space using eCLIP data and Chrom3D pointed to distinct splicing regulatory networks associated with peripheral chromatin (integrated within LADs) vs. centrally located one (*32*). Our list of proteins contains splicing machinery members allocated centrally, like FUS, AQR or DDX42, GEMIN5, as well as one of the classified peripheral active splicing factors (GPKOW). This potentially indicates that there might be a coordination between transcription and splicing at the pore. To pinpoint the location of these interactions in the 3D nuclear space, we have performed in situ proximity ligation assays (PLA), which allowed us to interrogate the spatial proximity and location of proteins of interest in Jurkat cells, whilst advantageously using antibodies against endogenous proteins in detriment of overexpression scenarios. Moreover, PLA allows the detection of spatially close proteins, within 40nm distance of each other.

**Figure 2:**
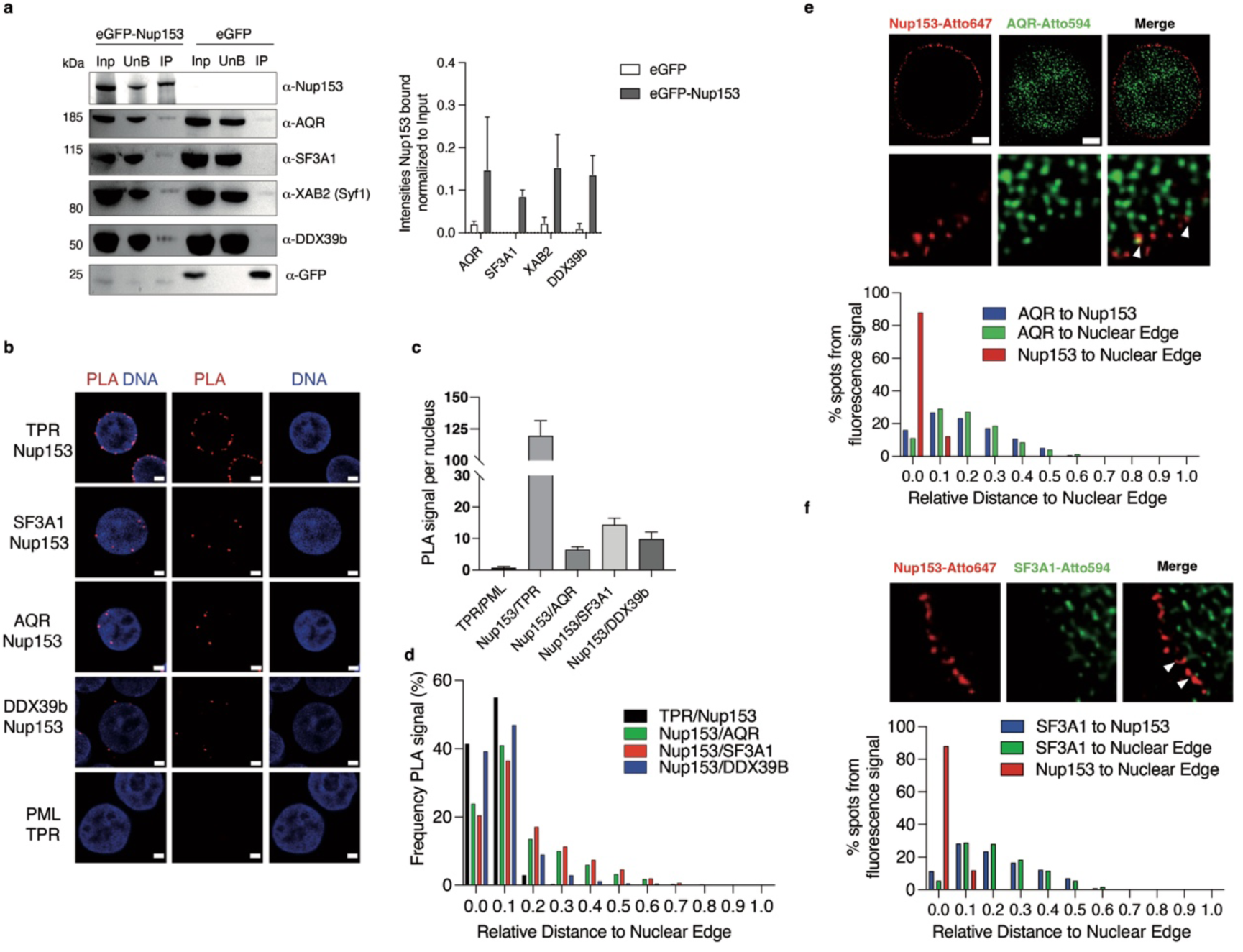
Splicing proteins associate with Nup153 at the level of the NPC: **a**. Co-IP of eGFP-Nup153 or control eGFP with spliceosome components in HEK293T cells, with quantification of n=3 experiments on the right. **b**. The *in-situ* PLA confirms an interaction of SF3A1, AQR and DDX39b with Nup153 at the nuclear periphery. Scale bar = 2 µm. **c**. PLA spots were normalized to the number of nuclei per field of view, plotted in bars representing the mean number of spots of ten fields of view. **d**. Nuclei and PLA signals were segmented, nuclear surfaces were created and the closest distance for each PLA spot to the nuclear surface was calculated in n > 200 number of nuclei. The distance was normalized to mean nuclear volume to obtain relative values, which were binned (bin size = 0.1) and plotted in a frequency distribution. STED microscopy resolves a peripheral portion of **e.** AQR or **f**. SF3A1 proximal to Nup153 in Jurkat cells, stained for Nup153-Atto-647 and AQR-Atto-594 (e) or SF3A1-Atto-594 (f). Scale bar = 2 µm. STED signals were transformed into spots and relative distances towards the Nup153 staining, or the nuclear edge, demarked by Hoechst staining (not shown), were obtained from n = 10 (AQR) and n = 9 (SF3A1) number of nuclei, were binned, and plotted in a histogram.

We observed PLA signals between Nup153 and its neighbor TPR, serving as a positive control, as well as between Nup153 and SF3A1, AQR, and DDX39b (**Fig.2b**). An interaction between TPR and PML was found negligible and served as a negative control (**Fig.2b**). The number of spots observed by PLA between Nup153 and SF3A1 was between 10-15% of the ones observed in the positive control Nup153/TPR, whereas DDX39b and AQR scored between 10 and 7 % of the Nup153 and TPR respectively (**Fig.2c**). We confirmed the spatial proximity between Nup153 and the IBC proteins AQR and XAB2 also in HEK293T cells, where we obtained an average of 8 and 12% of PLA spots per nucleus. B23 nucleophosmin, present mainly in the nucleolus, served as a negative control (**Fig.S2b-c**).

The positive control Nup153/TPR showed an entirety of 80% of signals at the nuclear periphery within 50 nm of the outer shell of the nuclei (**Fig.2d**). Consistently, the Nup153 and AQR, SF3A1 or DDX39b are spatially proximal and located at the similar (70nm) distance from the outer shells, therefore pointing to a peripheral interaction between Nup153 and the splicing machinery assessed here, in detriment of a nucleoplasmic interaction. A similar trend of spatial proximity between Nup153 and IBC proteins AQR and XAB2 at the nuclear periphery was also detected in HEK293T cells (**Fig.S2d**), suggesting that the association is conserved between different cell types, and confirming further our BioID data. We further classified this peripheral NPC-splicing proximity by using another Nup, Nup98. Nup98 is more internally located within the NPC than the basket Nup153 (*3*) . Nup98 has been shown to associate with DNA in drosophila and humans both in the nucleoplasm (*33–35*) and at the NPC (*10*, *35*) and through nucleoplasmic binding to an RNA helicase DHX9, implicated also in RNA processing (*36*). The PLA assay confirmed spatial proximity between Nup98 and AQR, SF3A1 and DDX39b (**Fig. S2e-f**). However, we detected a lower number of interacting spots between Nup98/DDX39b than with its Nup153 counterpart; a similar trend is followed between Nup98 and SF3A1 (**Fig. S2g**), suggesting that the interaction between splicing proteins and Nup98 might be further apart within the NPC structure. The proximity between Nup98 and the splicing proteins was still found at the peripheral compartment but, interestingly, there was a more widespread distribution interactions into the nucleoplasm, despite Nup98 being above Nup153 within the basket. We postulate that this could be ascribed to the dynamic nature of Nup98 associated with a possible nucleoplasmic pool or to a different antibody-specificity in comparison to Nup153 (**Fig. S2f**). Altogether, this data suggests that a pool of splicing proteins is at the nuclear periphery and, hence, available for contact with the NPC.

To investigate this further we have used the Stimulated Emission Depletion (STED) super resolution microscopy in Jurkat cells. Again, we found a peripheral pool of splicing proteins (**Fig.2e** and **2f**). The proportion of peripheral signals at the nuclear edge (0) is similar to that observed with the PLA counts (**Fig.2c and 2d**), indicating that a subset of pores accommodate splicing proteins in their proximity. In the Jurkat nuclei estimated to have a radius (4.7 +/-0.4 µm), AQR and SF3A1 signals were located in a distance range of 0.5 µm to 2.8 µm to the nuclear edge, i.e.in the 60-percentile of the nucleus. Thus, the nuclear center seems less enriched for AQR and SF3A1 which could be due to the presence of large sub-compartments in the nucleus center, like nucleoli. (**Fig.2e** and **2f**). Interestingly, in Jurkat cells in the absence of activation stimuli, AQR has focal (punctual) distribution throughout the nucleoplasm, possibly pertaining to its wide nuclear functions as an RNA helicase in the dissolution of DNA-RNA hybrids (R-loops) and DNA damage (*37*, *38*). Similarly, we do not observe SF3A1 assemblies in speckle-like compartments, in line with the proposed presence of RNAs throughout the nucleus (*28*, *29*).

### NPC association with IBC is splicing dependent

We investigated next if Nup153 could harbor splicing machinery assembling de-novo directly at the NPC. For that, we used PladienolideB (PladB), which stalls spliceosome assembly after the A complex formation (*39*). HEK293T cells were treated with PladB for 4h without massively impacting on cell viability (**Fig.S3a**). Analysis of intronic regions across a panel of reference genes confirmed a successful PladB treatment in blocking de-novo splicing machinery and inducing intron retention as previously shown (**Fig.S3b-e**). Pulling-down of transfected eGFP:Nup153 from HEK293T cells upon PladB treatment showed a decreased association between Nup153 and IBC members, AQR and XAB2, as well as DDX39b and SF3A1 (**Fig.3a**). Similarly, the pull-down of transfected Histidine-tagged AQR also yielded a lower detection of co-transfected eGFP:Nup153 in the presence of PladB. In the same experimental setting, blocking transcription with DRB (5,6-Dichloro-1-b-D - ribofuranosylbenzimidazole) (*40*), transcription elongation inhibitor, did not affect the interaction (**Fig.S3f**). Under PladB treatment conditions, the PLA using Nup153 and AQR or XAB2 antibodies against endogenous proteins reciprocated co-IP experiments, as we observed a reduction of PLA spots between untreated and PladB-treated HEK293T cells (**Fig.S3g**). Overall, this suggests that the association of the splicing proteins at the NPC is dependent on ongoing splicing.

**Figure 3:**
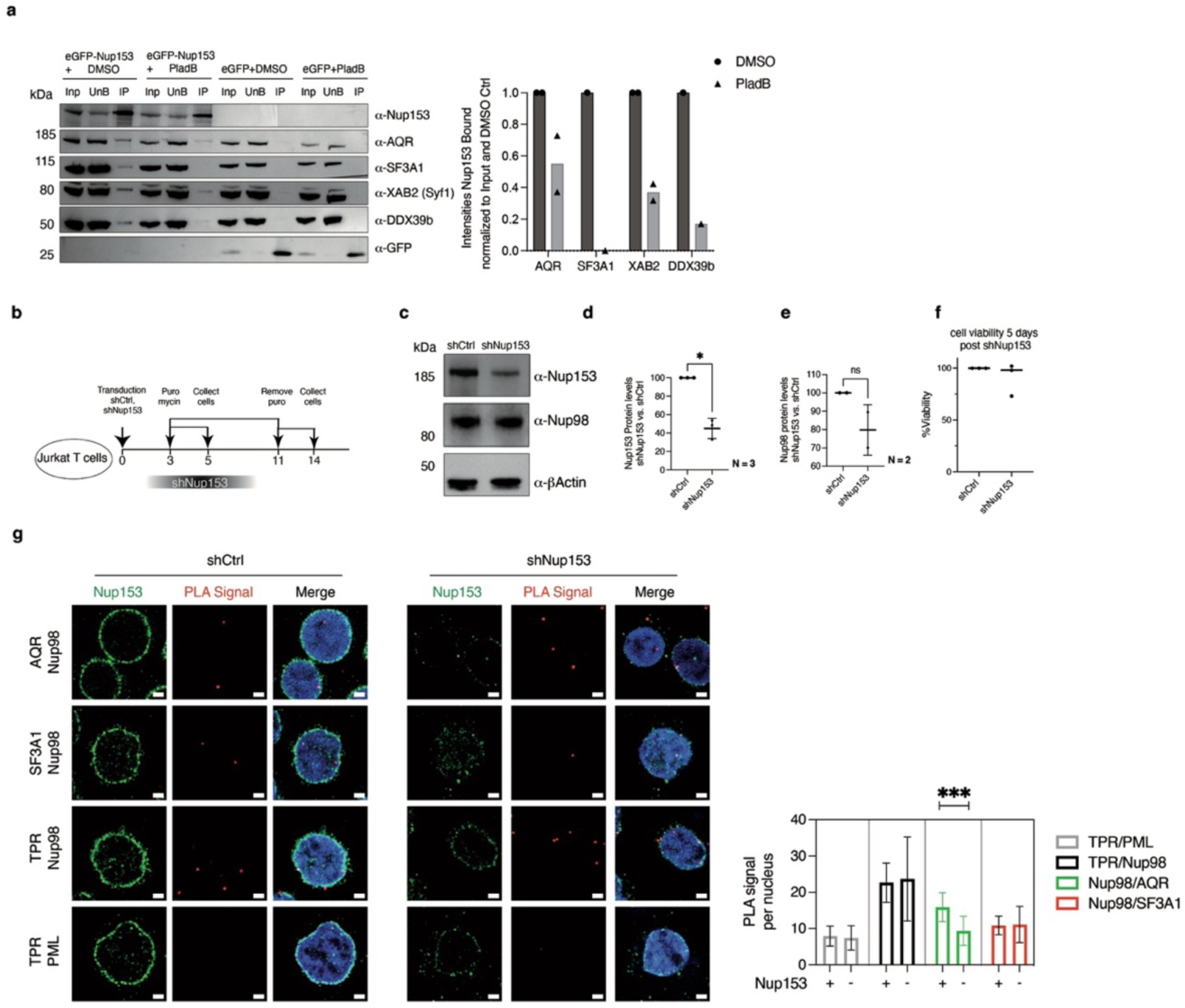
Nup153 aids the loading of spliceosomal proteins to the NPC. **a.** Co-IP for GFP-Nup153 or GFP alone from HEK293T cells transfected with eGFP-Nup153 or eGFP and treated with PladB or DMSO, followed by immunoblot for GFP, AQR, SF3A1, XAB2 and DDX39b. The graph on the right shows the intensities of bound fraction normalised to input and DMSO control. Bars present the normalized intensities for SF3A1 and DDX39b, and the means of two experiments for AQR and XAB2. **b**. Scheme of the experimental setup for the lentiviral transduction of Jurkat cells. **c**. Nup153 and Nup98 protein levels upon shNup153 depletion with representative immunoblot. **d**. quantifications of Nup153 (n=3) or **e**. Nup98 (n=2) protein levels, which were compared using unpaired t-test (* p < 0.05). **f**. Cellular viability, measured by the MTT assay at day 5 post shNup153 transduction. **g**. Exemplary PLA images of Jurkat cells transduced with shCtrl or shNup153 lentiviral vectors. In shNup153 transduced samples the cells were further stratified to Nup153 (-), where loss of Nup153 signal was detected, or Nup153 (+), where Nup153 staining of the nuclear rim seemed almost normal. PLA signal per nucleus quantification in Nup153 (+) (Tpr/PML: 95, Nup98/Tpr: 110, Nup98/AQR: 139, Nup98/SF3A1: 54 number of nuclei) vs Nup153 (-) (Tpr/PML: 99, Nup98/Tpr: 61, Nup98/AQR: 94, Nup98/SF3A1: 121 number of nuclei) cells of shNup153 transduced Jurkats cells. Mean number of PLA signals per nuclei were compared between shNup153 (+) and shNup153 (-) using unpaired t-test. *** p < 0.001.

### Nup153 aids the loading of splicing machinery

Having established the association between Nup153 and members of splicing machinery, we went on to understand if Nup153 plays a role in harboring splicing machinery.

We used shNup153 to knockdown Nup153 levels in Jurkat cells, compatible with the preserved nucleocytoplasmic trafficking at the NPC (*15*). Cells were under puromycin selection from day 3 and collected either on day 5 (**Fig.3b**) or left longer for the Nup153 recovery until day 11 with puromycin, and then collected at day 14 (shown in **Fig.5**). Cells on day 5 yielded the lowest levels of Nup153 (**Fig.3c** and **3d**) whilst preserving Nup98 protein levels (**Fig.3e**); cell viability was maintained, as determined by the cell viability MTT assay (**Fig.3f**).

We used the PLA assay to understand if, when Nup153 is depleted, the spatial proximity of the splicing proteins with the Nups can be altered at the level of the NPC in the nuclear periphery. For that, we used the Nup98 as a proxy for NPC interactions. We observe that the number of PLA spots between Nup98 and TPR is lower than between Nup153 and TPR (**Fig.3g)**, underlying the fact that Nup98 and TPR are not immediately adjacent, and possibly reflecting differences in antibody specificity. In shNup153 transduced cells, we could distinguish two cell populations, one where Nup153 could still be detected (marked as Nup153 (+) cells) and the other where Nup153 was affected (Nup153 (-) cells). We quantified the PLA signals between Nup98 and AQR or SF3A1 in both cell populations and observed statistically fewer interactions between Nup98 and AQR in Nup153 (-) cells, whereas this statistical significance was not observed with SF3A1 **(Fig.3g**). Importantly, the PLA spots detected between TPR and Nup98 (positive control) remained constant, suggesting that no major disruption in NPC structure occurred during the knockdown of Nup153; similarly, the interactions depicted by the negative control pair TPR/PML also remained invariant (**Fig.3g)**. This data suggests that Nup153 could potentially be involved in the recruitment of the splicing machinery. However, since Nup153 loss did not affect the broad recruitment of splicing proteins to the NPC, we cannot exclude other peripheral players acting as scaffolds of the peripheral pool of splicing proteins/machinery.

### Majority of Nup153 bound genes are proximal to LADs and expressed

To begin exploring the functional relevance of the NPC-splicing pool we employed DamID to retrieve genes binding to Nup153. We transduced Jurkat cells with lentiviral vectors expressing bacterial deoxyadenosine methylase (Dam)–Nup153 fusions to uniquely methylate DNA proximal to the Nup153 (*41*, *42*). To control for local variation in chromatin accessibility, parallel experiments expressed Dam methylase alone (Dam-only). The methylated DNA was then isolated and identified by sequencing. We stringently screened peripheral peaks by imposing a cutoff of proximity towards LADs borders (<100Kb) (*42*), hence allowing us to exclude Nup153 nucleoplasmic peaks that would otherwise hinder subsequent analysis. A total of 1627 peaks were identified, distributed in all chromosomes, according to the distribution of the number of peaks per megabase of each chromosome (**Fig.4a**). Identified peaks were found in distal intergenic regions in 65% of cases (**Fig.4b**), while the rest (700 total peaks) were found in genic regions, with intronic regions (28% of peaks) being the prevalent sites of Nup153 binding (**Fig.4b**). In total, we identified 510 Nup153 peaks associated with 461 protein-coding genes (**Fig.4c**). By using Jurkat Lamin_DamID data (*42*) we calculated the genomic distance of Nup153-bound genomic regions (i.e. their respective DamID peaks) to the closest LAD border (**Fig.S 4d**). Unlike randomly generated peaks, DamID:Nup153 peaks were indeed accumulated near the LAD border with a much narrower distribution, especially noticeable for genic Nup153 peaks (**Fig.S4e**). Because of the stringency criteria we applied to define DamID: Nup153 signals, we conclude that they correspond to the peripheral (and NPC bound) pool of Nup153, rather than to the nucleoplasmic Nup153.

**Figure 4:**
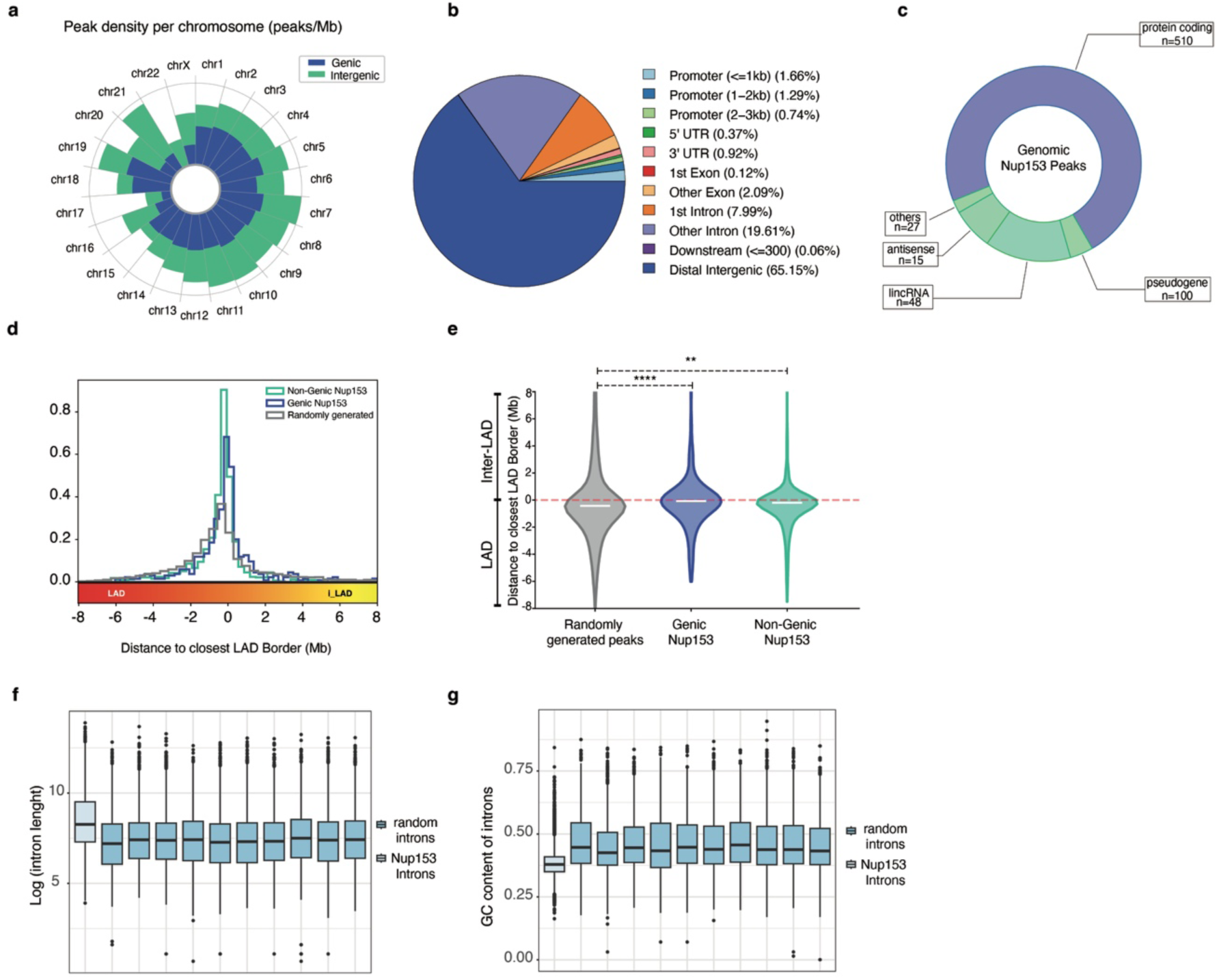
Mapping of Nup153 associated genic regions with DamID reveals a group of peripheral long genes with multiple introns and exons. **a.** Distribution of Nup153 genic and non-genic peaks per chromosome. **b.** Genomic features of Nup153 peaks. **c.** Features of Nup153 genic peaks. **d.** Distribution of Nup153 genic and non-genic peaks as well as a population of randomly similarly sized genes in relation to LAD borders. **e** Average distribution of Nup153 genic and non-genic peaks at the LAD border in comparison to a population of randomly similarly sized genes **f.** Average length of introns in the Nup153 genes in comparison to ten random populations of equally sized numbers of genes. **g.** GC content of Nup153 genes in comparison to ten random populations of equally sized numbers of genes.

**Figure 5:**
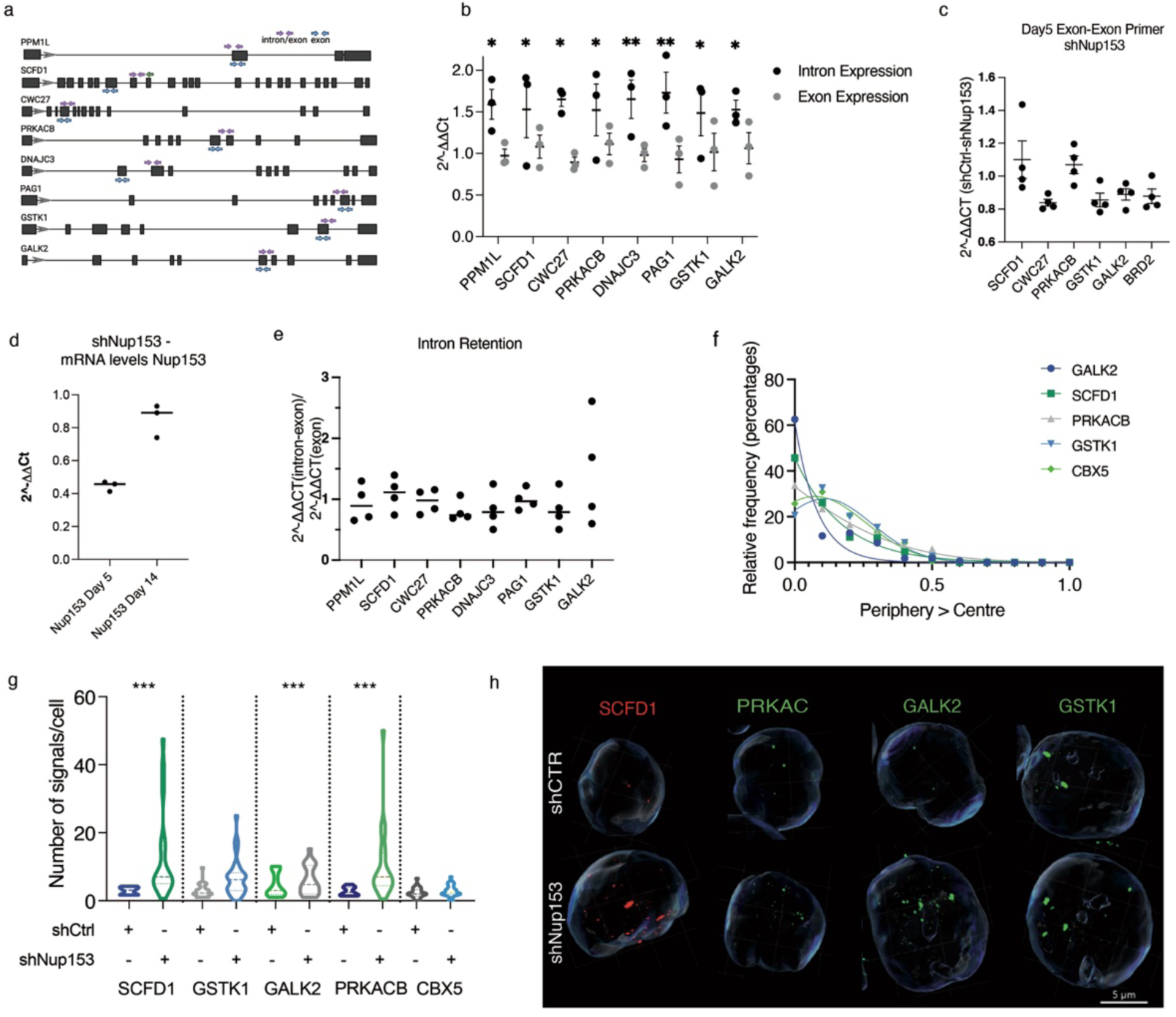
Nup153 contributes to the efficient splicing of Nup153 bound genes. **a**. Scheme of primers designed in a sub-set of eight Nup153 bound genes. Exon primer pairs (blue) differ from intron-exon primer pairs (pink). **b**. qPCR of shCtrl and shNup153 samples extracted 5 days post lentiviral knockdown, with exon and intron-exon primers described in (**a)**. TBP served as a normaliser, with Intronic-exonic primer pairs all showing statistically significant change in shNup153 samples when compared shCtrl. **c**. qPCR of shCtrl/shNup153 transduced Jurkat cells using exon-exon primers of a sub-set of genes including TBP as a normaliser. **d**. qPCR of Nup153 expression levels at day 5 or day 14 post transduction using TBP as a normaliser. **e.** qPCR of shCtrl and shNup153 samples extracted 14 days post lentiviral knockdown was performed using exon and intron-exon primers described in a. **f.** Relative distance of frequency of SABER-FISH of shCtrl and shNup153 samples 5 days post transduction in a panel of genes. **g.** Number of SABER-FISH signals per cell in shCtrl and shNup153 samples 5 days post transduction in a panel of genes. **h.** Example images of SABER-FISH in shCtrl and shNup153 samples 5 days post transduction in a panel of genes. Experiments were performed in triplicate and qPCR data is shown in mean and include standard deviation. Multiple T-test was (*** p < 0.001) .

However, we observed that as opposed to mainly repressed genes in LADs, Nup153 genes are largely transcribing, as emerging from comparisons with cognate Jurkat RNA seq datasets (**Fig.S4a**). Out of 461 Nup153 genes, 211 were not expressed, while others were either mid to low expressed (235 genes) or high expressed (15 genes). These expression profiles of Nup153 bound genes differ from genes mapped in LADs, where 80% of them are found repressed, and have a closer resemblance to a randomly selected group of protein-coding genes. We next analyzed chromatin patterns on Nup153 genes and compared them with a similar group of randomly sized population of protein-coding genes or with LAD genes. We found that Nup-bound genes represent an intermediate group harboring H3K4me3 positive histone marks at similar abundance as protein-coding genes (**Fig.S4b**), contrary to LADs. When compared to LADs, Nup153 bound genes seem to have higher occupancy of positive histone modifications and RNAPII, while at the same time, their H3K27me3 occupancy is similar to LADs (**Fig.S4b**). While previous binding profiles of nuclear basket nucleoporins indicated their association with super-enhancers, DNA elements enriched in H3K4me1, H3K27ac, BRD4, and Med1 (*12–14*), we could not observe a consistent association of Nup153 peaks with regions defined as SE in the cognate cell line (not shown). It is possible that the heterogeneity in chromatin marks decorating the DNA bound by Nups (Nup153) could be hindered by the use of bulk genomic analysis.

### Nup153-associated genes are enriched in long introns with low GC content

We next characterized Nup153 bound genes enquiring whether they could benefit from the assembly of splicing machinery at the NPC level. From the total identified 461 protein coding genes with 510 Nup153 peaks associated with them (**Fig.4c**) we began investigating their intron profiles and GC content. The initial step of spliceosomal assembly across exons or introns occurs respectively via exon or intron definition mechanisms. Long introns with lower-than-average GC content are a hallmark of the so-called intronic definition of splicing (*43*) . Recently, owing to the GP-Seq, which maps the radial position of genes based on their GC content (*44*), it was proposed that genes with longer than average introns and lower than average GC content localize mainly at the nuclear periphery (*32*)

We therefore looked at the characteristics of Nup153 introns by assessing their length and overall GC-content. We observed that Nup153 introns are significantly longer than introns from 10 control groups of 500 averagely selected genes (**Fig.4f**). Concordantly, the GC-content of Nup153 introns was markedly lower than that of control genes (**Fig.4g**).

To assess if our Nup153_DamID mapped regions are indeed localized in the periphery, we aligned them with Lamin_DamID data from Jurkat cells and averaged them based on their GP-Seq scores from HAP1 cells (*44*). A large portion of Nup153 genes mapped in the outermost shells (periphery) of the nuclei (**Fig.S5a)**, consistent with the notion that GC-low genes are preferentially found in the periphery. Visual representation across a panel of chromosomes (**Fig.S5b**) shows the peripheral location of a pool of Nup153 genes, a subset that was depicted for use in subsequent analysis.

### Nup153 associated genes are aberrantly spliced in the absence of this nucleoporin

Equipped with a group of genes bound to Nup153 we sought to assess the splicing of these genes in the peripheral nuclear compartment. To begin assessing the possible implications of shNup153 on splicing, we have designed two sets of primers, one set covering the intron and the exon junction of selected introns and a second set on the most proximal exon (scheme in **Fig.5a**). Using the exonic primers we could observe by real-time qPCR that exon amplification does not change significantly in shNup153 conditions with respect to the shCtrl (**Fig. 5b**). However, the intronic amplification resulted higher in shNup153 conditions, indicative of the introns that were still present upon Nup153 knockdown. Albeit small, these differences are statistically significant for the tested Nup153 bound genes, and importantly this trend was not observed for a non-Nup153 bound gene (**Fig.S6a**). We additionally tested some of the genes by designing primers across exons relatively similar in size flanking a short intron, compatible with an accurate PCR amplification (**Fig.S6b**). We were able to observe an additional, intronic (upper) band in Nup153 knockdown conditions for PRKACB, SCFD1 GSTK1and GALK2 genes, whereas only one band, pertaining to the spliced (exon-exon junction) amplicon, was detected in shCtrl control condition (**Fig.S6c**). Consistently, when using these primers in qPCR, we could observe that the amplification was slightly diminished for some of these genes suggesting less availability of the spliced form, statistically significant for GALK2 (p=0.049), GSTK1 (p=0.04) and CWC27 (p=0.004) (**Fig. 5c**), suggesting that the presence of the long intron in between the two exons hindered qPCR amplification (typically optimal for short amplicons).

Following a Nup153 recovery, which we observed after 14 days in culture (**Fig. 5d**), we assessed if intron retention could be recovered once Nup153 levels are restored. Indeed, we observed that, for most of the assessed genes, intron retention is no longer observed as Nup153 levels recover (**Fig. 5e**). Altogether this data suggests that Nup153 could potentially have an effect on intronic retention of the genes to which it is bound. Despite the fact that the techniques used here did not favor the unstable nature of introns, nor had the required sensitivity to detect this in great detail, we conclude that the Nup153 presence is required for the recruitment of splicing proteins with functional implications on the subset of Nup153 bound genes.

We next employed signal amplification by exchange reaction fluorescence in situ hybridization (SABER-FISH) (*45*) using intronic probes to detect RNA of SCFD1, PRKACB, GSTK1 and GALK2. This method amplifies signal intensity and therefore provides a good basis for inspection of low/transient transcript forms. We were able to visualize the presence of a site of transcription where accumulation of these probes was close to the periphery for all except for GSTK1, which showed a wider nuclear distribution, similar to CBX5 control region not bound by Nup153 (**Fig.5f**). The observed distribution of Nup153 bound genes seems consistent with results of the cross-sectioning of our Nup153 DamID and GP-Seq data (**Fig.S5b**). We complemented SABER FISH with IF in order to count intronic FISH signals on cells depleted from Nup153.

We detect a higher number of FISH signals in Nup153 depleted cells in comparison to shCtrl, indicating the presence of intronic probes in the absence of Nup153 (**Fig.5f, g** and **5h**). This was statistically significant for SCFD1, GSTK1, PRKACB, and a similar trend was followed by GALK2; importantly FISH signals for our negative control CBX5 were not altered upon Nup153 knockdown. Together, this suggests that Nup153 has an effect on splicing of the genes that are transcribing within its vicinity.

## Discussion

In this study we used BioID to systematically address the interactome of basket Nups TPR and Nup153. Apart from the expected interactors, pertaining to transcription, DNA replication or trafficking functions, we report the presence of splicing proteins belonging to all spliceosomal complexes. The relevance of our finding is reflected in the fact that an abundant number and a wide range of splicing proteins were identified interacting with NB. In support, nuclear transporter receptors binding to NPC yielded very few and non-statistically significant interactions with spliceosomal protein (*30*), suggesting that transport Nups less likely interact with splicing machinery. Our findings align with the enrichment of RNA splicing and metabolism proteins observed by SILAC approaches targeting TPR binding factors pinpointing to an enrichment of RNA splicing and metabolism (*46*), supported by a previous proteomics study, which identified mRNA processing, splicing and export proteins in the TPR interactome (*16*). Moreover, a reciprocal BioID analysis of the XAB2 component of the Intron Binding Complex, belonging to the B* active splicing complex, in mice liver identified several nucleoporins, including Nup153 (*47*), further strengthening the notion that different splicing components can be found proximal to basket nucleoporins. The splicing proteins identified here were also characterized in the extensive splicing factor network either in the peripheral (hnRNPA, GPKOW, PPIE) or in the central splicing factor (FUS, DDX42, GEMIN5l, AQR) subnetwork (*32*). Altogether, these support the role of the NPC as a platform coordinating transcription mRNA splicing and export. Furthermore, our study suggests that splicing machinery is associated with the NPC across two very disparate cell lines. A protein proximity interactome study of the recently described NPC basket component guanylate-binding protein (GBP)-like GTPase (GBPL3) in Arabidopsis also identified RNA splicing and processing proteins enriched at the level of the NPC basket (*48*). This implies that the splicing machinery’s physical and most likely functional association with the NPC basket represents a conserved feature across various species.

Taking the PLA spot counts at face value, we observe splicing proteins to be located on only 10% of pores, highlighting the heterogeneous nature of NPCs. This opens an important question regarding the number of pores effectively associated with transcription and import/export functions, and supports the idea of varied protein content at the NPC exerting different non-canonical NPC functions (*1*, *6*). In fact, the heterogenous nature of protein composition at the NPC could account for the polycomb repressive complex (PRC1) associated with Nup153 and the gene inactivity reported in embryonic stem cells (*9*) contrasting the spatial relationship between Nup153 and Super Enhancers (SE) (*12–14*, *49*, *50*). Approximately ∼10% of SE are located at the NPC, forming phase-separated compartments that govern positive transcription of a specific subset of genes (*14*). Whether splicing components congregate in specific pores through phase-separation prompts future investigations.

The functional significance of NPC-splicing associations began to emerge when we mapped the Nup153 bound genome by DamID, and identified 461 genes, which according to their proximity to LAD borders (*42*) and based on GP-Seq data intersection (*44*) seem to be mostly peripheral. The intertwining between nuclear pore complexes and lamins have been revealed by Cryo-ET (*51–53*). Accordingly, a recent work has also placed Nup153 as a central player in the recruitment of LaminB1 (*54*), justifying the close ties between these two proteins. and suggesting that Dam would likely methylate adenines on GATC that are spatially converging due to the proximity between Nups and adjacent LADs.

The Nup153 genes resulted in having longer than average introns and lower than average GC content. Genes with long introns and low GC content were previously shown to be enriched at nuclear lamina (*55*). However, Nup153 genes do not pertain to the categories of genes described to be LMNA associated (microtubule organization, ncRNA processing), suggesting that they represent a different portion of peripheral genes. Interestingly, the length of genes and number of introns seem to correlate with their distance to speckles. The sub-categorized transcripts and their relation to speckles could be defined as stably or transiently enriched at speckles (Groups A and B, respectively), or speckle-distant transcripts, characterized with long introns and low GC content (C group). These features, could well correspond to the Nup153 associated genes, as speckle-distant group of spliced genes (*26*, *27*), and will be further explored in the light of the widespread availability of pre-mRNA throughout the nucleoplasm (*28*, *29*) as well as in relation to the role of GC content in defining the gradient of splicing proteins in different cell types.

Cell type specific differences in splicing patterns might further be influenced by the activation status of cells, particularly relevant for the immune cells. Previously reported to play a role in detachment of certain genes from nuclear lamina (*42*), cell activation status of lymphoid or myeloid lineages might also influence the extent of Nup153 – genome attachment. Indeed, we have previously observed that during T cell activation, certain genes are moved towards the nuclear periphery, and possibly towards the NPC (*56*), indicating that T cell activation might also change the profile of NPC associated genes.

A progressive increase in intron length during evolution has resulted in the human genome having significantly longer introns than other mammals (*57*). Human cells therefore had to evolve strategies to facilitate the identification of exons located between long introns, but it remains still unclear how spatially distant splice sites are brought into proximity when introns are thousands of nucleotides apart (*58*, *59*).

Our study suggests that Nup153 facilitates the recruitment of splicing proteins to long introns. Interestingly, the pentameric Intron Binding Complex (IBC) and its helicase component AQR (*60*), known also as the Intron Binding Protein 160 (IBP160), bind splice branch site(s) in intronic regions (*61*) to activate the spliceosome (*62*) and regulate numerous splicing events in mammalian cells (*22*). Thus, the binding of Nup153 to the underlying genome would enable the tethering of the splicing machinery to fully process these transcripts before their export.

Taken together this study supports the gene gating theory and the accumulation of evidence pointing towards the presence of transcription and export, and now, splicing at the NPC level.

### Limitations of the study

This study points to the presence of splicing components at the NPC, which in tandem with transcription and export would pose valuable support for the gating of induced genes moving towards the periphery during stimuli/induction. In light of this, the experiments performed here in a steady-state scenario provide the grounds for further exploitation of the functional impact of these rounded machine networks at the NB in an inducible scenario. Whether Nup153 harbours these three complex machineries through phase separation, similar to SE compartmentalization, remains to be understood. In the case of Nup153-SE, interactions are mediated through the C-terminal FG-rich Intrinsically Disordered Region (IDR) of Nup153, whereas the central Zn finger containing domain of Nup153 is responsible for attachment of Nup153 to DNA (*63*). Future investigations can explore if the FG-Nup153 IDR tethers these components to the periphery.

## Data Availability

All relevant data supporting the key findings of this study are available within the article and its Supplementary Information files or from the corresponding authos. The DRIPc from activated CD4+ T cells is available under can https://github.com/bsrezovic/Nup153

## Code availability

All code accompanying this paper is available at https://github.com/bsrezovic/Nup153

## Acknowledgments

We would like to acknowledge the Infectious Diseases Imaging Platform (IDIP) platform for microscopy support, as well as the Deep L Genomics Core Facility Heidelberg and EMBL genomics facility and c. ATG Tübingen for their technical support.

We would like to acknowledge Exaltum, LTD for their excellent bioinformatic support services.

## Author contributions

ML and IC designed the experiments; IC, LS, CPG, AU, MRV, TS, GT, JH, EM performed and analyzed the experiments. IC and EM performed BioID and analyzed with TS BioID data, IC, LS, GT, VL performed and analyzed microscopy data, CPG, DG, JdlH, GS performed bioinformatic analysis of the DamID Nup153 data and all the LAD and chromatin comparison datasets. ML and IC wrote the manuscript. ML conceived the study and supervised the project. All of the authors commented on and edited the manuscript.

## Material and Methods

can be found in the Supplementary data file

