## Supplemental Figures and methods for "Nucleoporin Nup153 docks the splicing machinery to the nuclear pore for efficient mRNA processing"

8 Present address: College of Health, Medicine and Life Sciences, Division of Biosciences, Brunel University London, Uxbridge, UB8 3PH, U.K.
9 Present address: Department of Experimental Physics, Saarland University, Saarbrücken, Germany
10 Present address: Friedrich Miescher Institute for Biomedical Research, Fabrikstrasse 24, 4056 Basel, Switzerland

### Supplementary Figures and Figure legends

a

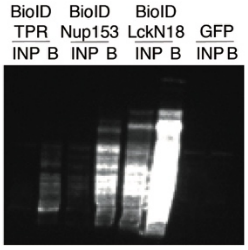


b

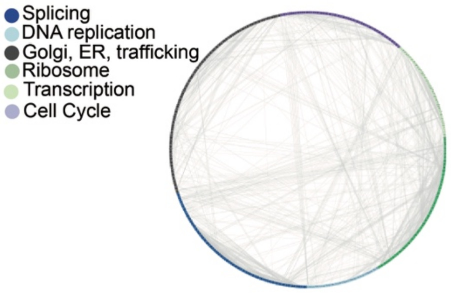


c

| Description | Spliceosomal | | | | |
|---|---|---|---|---|---|
| serine and arginine rich splicing factor 6(SRSF6) | E | A | B | B* | C |
| DExD-box helicase 39B(DDX39B) | E | | | | |
| catenin beta like 1(CTNNBL1) | C | P | | | |
| SLU7 homolog, splicing factor(SLU7) | C | P | | | |
| pleiotropic regulator 1(PLRG1) | C | P | | | |
| DEAH-box helicase 8(DHX8) | C | P | | | |
| chromosome X open reading frame 56(CXorf56) | C | P | | | |
| family with sequence similarity 50 member A(FAM50A) | C | P | | | |
| DEAH-box helicase 16(DHX16) | B* | C | | | |
| papillary renal cell carcinoma (translocation-associated)(PRCC) | B* | | | | |
| G-patch domain containing 1(GPATCH1) | B* | C | | | |
| FUS RNA binding protein(FUS) | B* | C | | | |
| G-patch domain and KOW motifs(GPKOW) | B* | C | P | | |
| CWC22 spliceosome associated protein homolog(CWC22) | B* | C | P | | |
| Sin3A associated protein 18(SAP18) | B* | C | | | |
| crooked neck pre-mRNA splicing factor 1(CRNKL1) | B | B* | C | P | |
| WW domain binding protein 11(WBP11) | B | | | | |
| pre-mRNA processing factor 3(PRPF3) | B | | | | |
| breast carcinoma amplified sequence 2(BCAS2) | B | B* | C | P | |
| CWC15 spliceosome associated protein homolog(CWC15) | B | B* | C | P | |
| DEAD-box helicase 23(DDX23) | B | | | | |
| tuftelin interacting protein 11(TFIP11) | B | | | | |
| ubiquitin specific peptidase 39(USP39) | B | | | | |
| peptidylprolyl isomerase H(PPIH) | B | | | | |
| small nuclear ribonucleoprotein U5 subunit 200(SNRNP200) | B | B* | C | P | |
| aquarius intron-binding spliceosomal factor(AQR) | B | B* | C | P | |
| peptidylprolyl isomerase E(PPIE) | B | B* | C | P | |
| ISY1 splicing factor homolog(ISY1) | B | B* | C | P | |
| XPA binding protein 2(XAB2) | B | B* | C | P | |
| splicing factor 3a subunit 1(SF3A1) | A | B | B* | C | P |
| splicing factor 3b subunit 6(SF3B6) | A | B | B* | C | P |
| SURP and G-patch domain containing 1(SUGP1) | A | | | | |
| splicing factor 3a subunit 3(SF3A3) | A | B | B* | C | P |
| small nuclear ribonucleoprotein polypeptide A(SNRPA) | A | B | | | |
| small nuclear ribonucleoprotein polypeptide A'(SNRPA1) | A | B | B* | C | P |
| small nuclear ribonucleoprotein polypeptide B2(SNRPB2) | A | B | B* | C | P |
| heterogeneous nuclear ribonucleoprotein A/B(HNRNPAB) | A | | | | |
| RNA binding motif protein 10(RBM10) | A | | | | |
| pre-mRNA processing factor 4(PRPF4) | | B | | | |

Fig. S1

**Fig. S1**: **BioID followed by mass spectrometry for Nups Tpr and Nup153 showing enrichment in splicing proteins across spliceosome. a.** Example of a Streptavidin immunoprecipitation on HEK293T cells transfected with BirA:Tpr, BirA:Nup153 and a positive control BirA:LckN18 and GFP (not fused to BirA, negative control). Proteins isolated using Streptavidin beads were on-bead trypsin digested for indirect identification; biotinylated beads were eluted in an additional step using a mixture of ACN and TFA for direct Mass Spec identification. Example of eluted samples ran on a a western blot using an antibody against Streptavidin-HRP. **b**. STRING network analysis of the five clusters seen in **Fig.1c** was exported to Cytoscape for visualization purposes, Circular Layout emphasizes group and tree structures within the network with a connectivity structure. **c.** The spliceosomal proteins depicted within our BioID are displayed across the several spliceosomal components (E, A, B, B*, C and P) and not restricted to one.

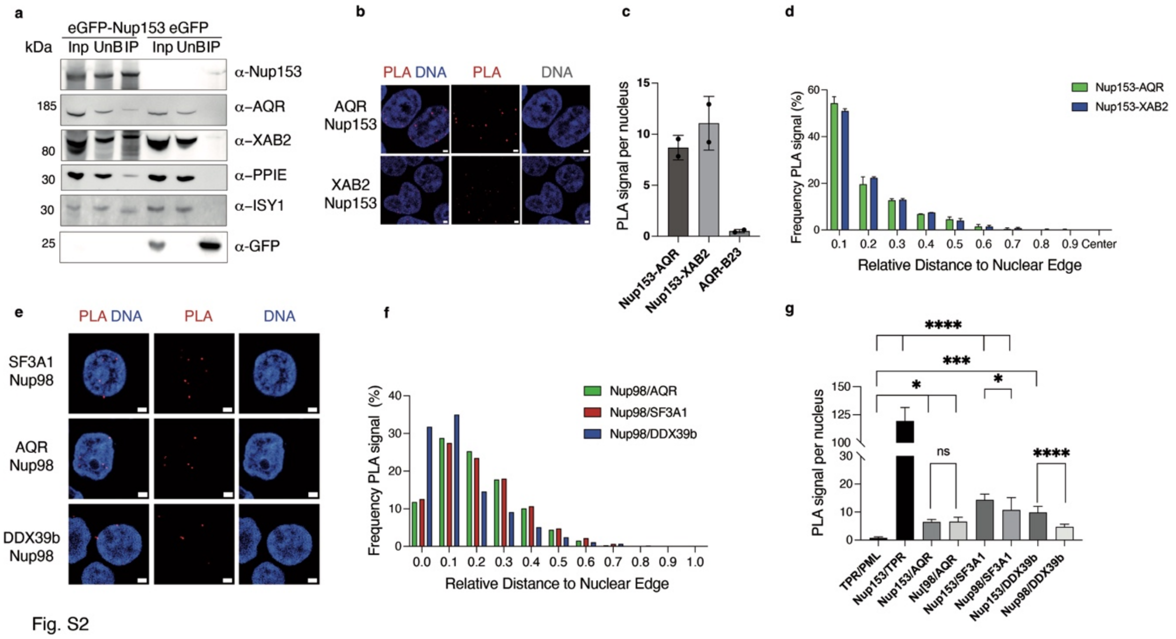


Fig. S2

**Fig.S2: Splicing proteins associate with nucleoporins Nup153 and Nup98 at the level of the NPC.** **a**. Co-IP of eGFP-Nup153 or eGFP with components of the IBC complex. **b**. PLA assay showing the spatial proximity between Nup153 and AQR/XAB2 of the IBC in HEK293T cells. **c**. Quantification of PLA signals per nucleus from HEK293T cells **d**. Relative distance of the frequency of PLA signals between Nup153 and AQR/XAB2 to nuclear envelope. **e**. PLA signals between Nup98 and splicing components SF3A1, AQR and DDX39b in Jurkat cells **f**. Relative distance of the frequency of PLA signals between Nup98 and AQR/SF3A1/DDX39b to the nuclear envelope. **g**. Quantification of the number of PLA spots per nucleus of interacting splicing components with Nups 153 and 98. Total nuclear PLA spots were normalized to the number of nuclei per field of view, which were then plotted in bars representing the mean number of spots of ten fields of view, which were compared using one-way ANOVA with Dunnett's multiple comparison test. PLA spots of splicing proteins with Nup153 or with Nup98 were compared using unpaired Welch's t-test. * $p < 0.05$; *** $p < 0.001$; **** $p < 0.0001$.

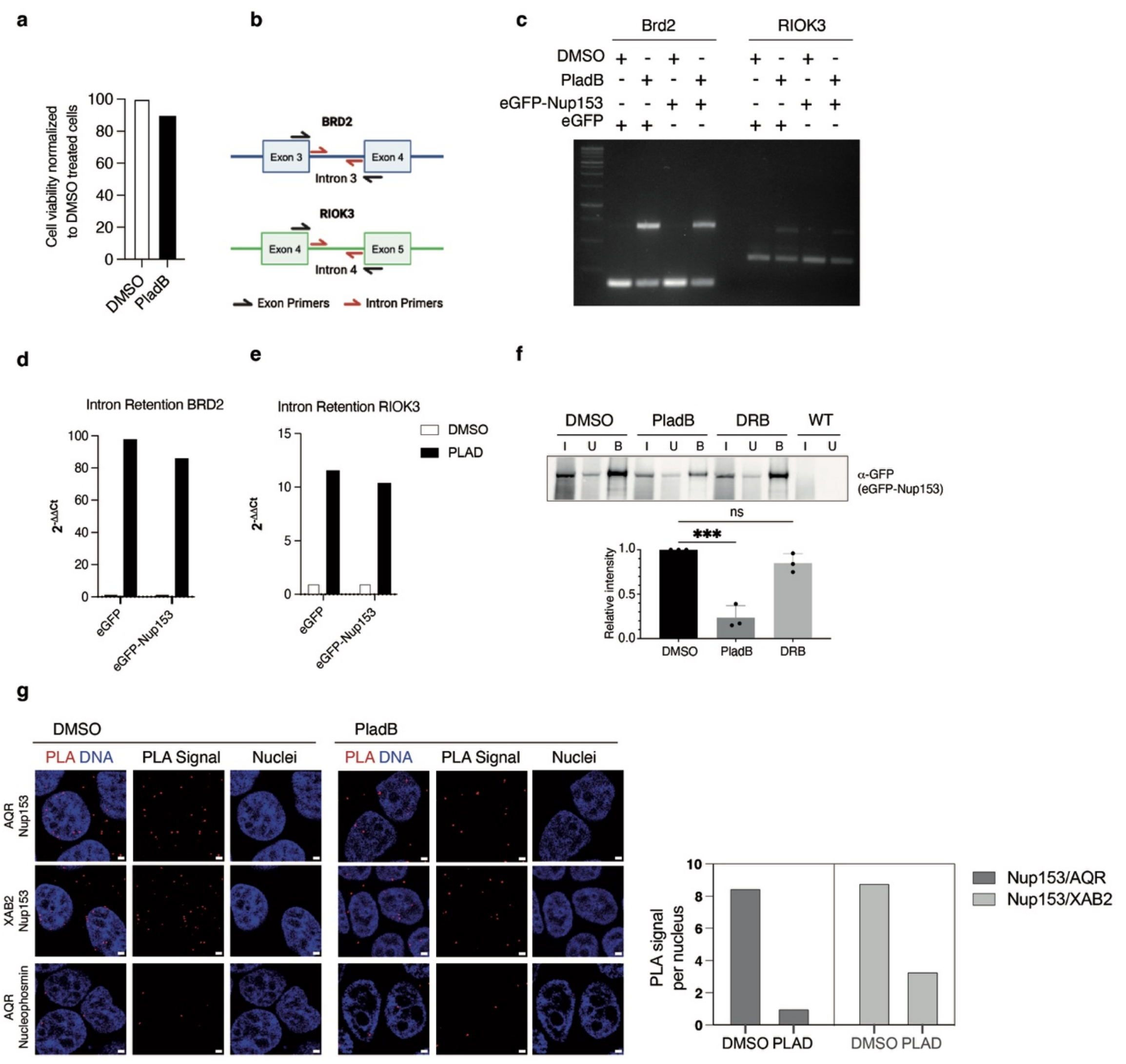


Fig. S3

**Fig.S3: Stalled spliceosome assembly affects Nup153 interactions with the splicing proteins. a.** Cell viability in PladB treated cells. **b.** Position of designed primers used to detect intron amplification. Exon primers were used for PCR, intron primers for qPCR. **c.** Amplification of Brd2 and RIOK3 control genes for PladB treatment using primers from 11, visualized with agarose gel electrophoresis **d.** qPCR of BRD2 using newly designed qPCR intron primers from b**. e.** qPCR of RIOK3 using newly designed qPCR intron primers from b. **f.** Co-IP between AQR-His-Flag and eGFP-Nup153 upon plasmid transfected into HEK293T cells and treatment with PladB or with DRB to halt transcriptional elongation. Immunoprecipitation was performed using Flag for AQR, and WB was Quantification of 3 experiments is shown in the lower panel. **h**. Representative images of the PLA assay performed in HEK293T cells upon PladB treatment; quantification of PLA signals obtained for interaction of Nup153 with IBC members AQR and XAB2 is shown on the right.

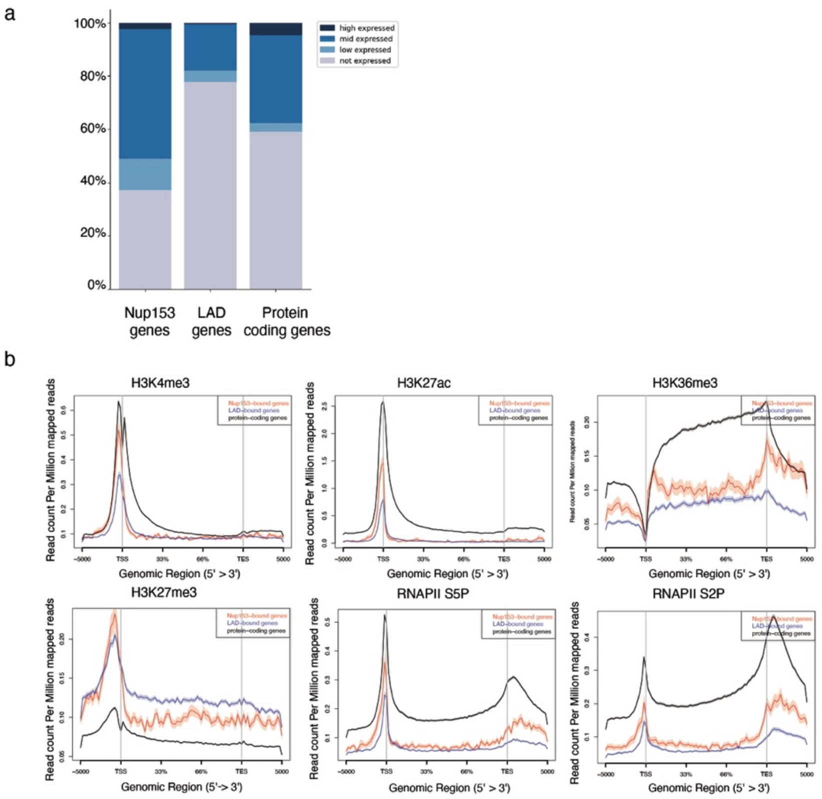


Fig. S4

**Fig.S4. Nup153 genes show an intermediate phenotype between LADs and protein coding genes .** **a.** Expression levels of 510 genes found within the Dam:Nup153 dataset in comparison to a similar-sized population of protein coding and LAD-bound genes from Robson et al (*1*). **b.** Average profiles of histone modifications and RNAPII across the coding region (TSS up to TES) of Nup153-bound and similar-sized LAD-bound and protein coding genes.

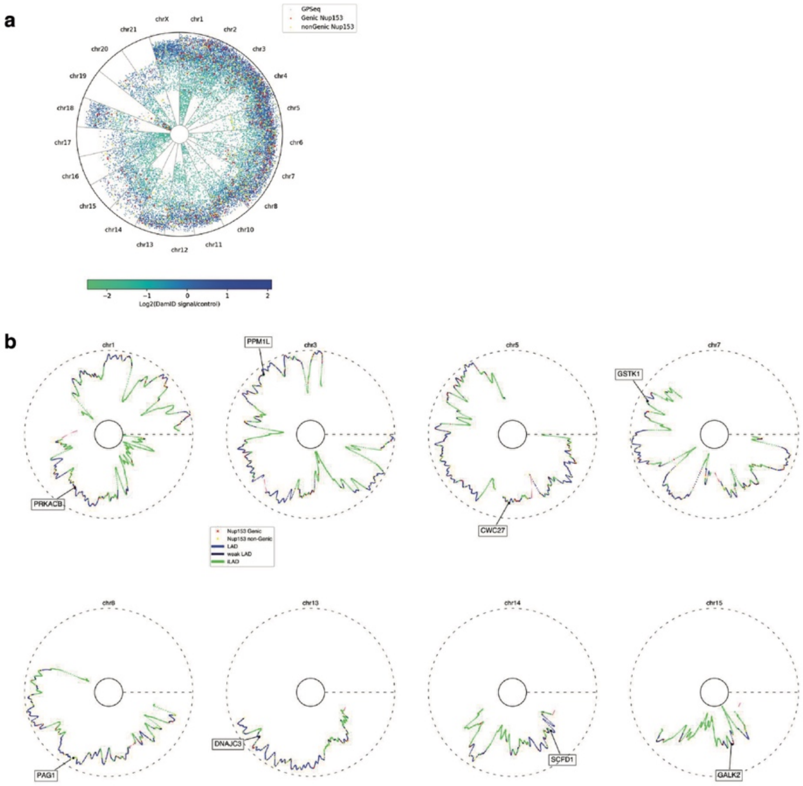


Fig. S5

**Fig.S5: Peripheral location of DamID:Nup153 peaks in relation to GP-seq.**

**a.** Radial distribution of GP-Seq, DamID:Nup153 genic and non-genic peaks across chromosomes from the center to the outmost peripheral shell. **b**. Panel of chromosomes depicting the radial distribution of GP-Seq, DamID:Nup153 genic and non-genic peaks including a panel of genes used in follow up analysis.

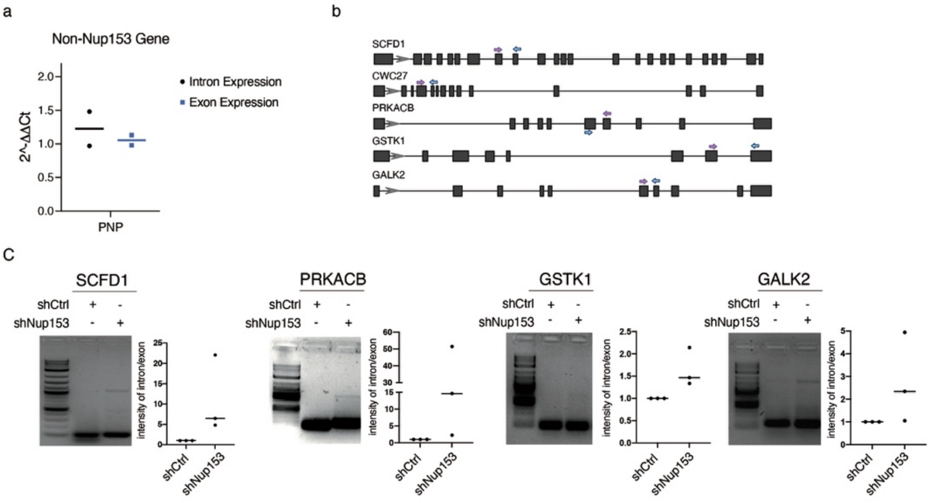


Fig.S6

**Fig.S6: Transcripts of Nup153 bound genes show increased intron retention in the absence of Nup153 a.** qPCR of shCtrl and shNUP153 samples extracted 5 days post lentiviral knockdown was performed using PNP exon and intron-exon primers; TBP was used as a normaliser. **b**. Schematics of primers designed in a sub-set of five Nup153 bound genes covering two exons intercalated by relative short primers compatible with PCR amplification. **c.** PCR of shCtrl and shNup153 samples extracted 5 days post lentiviral knockdown was performed using primers shown in b.

**Table S1:** List of proteins obtained in the BioID analysis

**Table S2**: List of antibodies used troughout the work in immunoflorescence, PLA and WB together with all primer sequences.

### Methods

#### Cell culture

Cells were grown at 37 °C and 5 % CO2 in a humidified incubation chamber. HEK293T cells (Pear et al. 1993) were maintained in Dulbecco's Modified Eagle's Medium (DMEM; Thermo Fisher Scientific) containing 4.5 g/L D-glucose and L-glutamine supplemented with 10 % fetal calf serum (FCS; Sigma-Aldrich, USA), 100 U/ml penicillin and 100 µg/mL streptomycin (PAN Biotech, Germany). T-lymphocyte Jurkat E6.1 (ATCC) or JTAG cells (E6.1 stably overexpressing SV40 Large T-antigen, kind gift from Fackler) were maintained in RPMI 1640 (Thermo Fisher Scientific) containing L-glutamine supplemented with 10 % FCS, 100 U/mL penicillin and 100 µg/mL streptomycin.

**cDNA vectors and shRNAs**

Construct for overexpressing p-EGFP-Nup153 (obtained through Euroscarf, Jan Ellenberg lab, used previously in (*2*)) was used in co-immunoprecipitations in tandem with pWPIeGFP (kind gift from Didier Trono). pENTER_AQR-HIS-FLAG was purchased from Vigene Biosciences. pLgwBirAmChDamNup153 or pLgwBirAmChDam (kind gifts from E.Lemke) were used in BioID and DamID experiments. BioID:LCKN18 (kind gift from O.T. Fackler) was used as a positive control for BioID experiments and to clone BioID:TPR by using BspEI and ApaI restriction enzymes on pCDHOST-TPR (kind gift from T.Misteli).

The shRNAs targeting NUP153 was constructed based on pLKO.1 lentiviral vector and targeting sequence was obtained from the publicly available TRC cloning portal of sigma (TRCN0000308266 CATTGGTGTTGTACTCAAT) and selected with puromycin.

**Transfections, Lentiviral transduction**

For co-immunoprecipitations, HEK293T were transfected with pEGFP-Nup153 or with a pWPIeGFP using JetPRIME (Polyplus) as instructed by the manufacturer's protocol. After 24 h, cells were either harvested directly for co-immunoprecipitation (Co-IP) or treated with 200 nM Pladienolide B (PLAD) for 4h (or 100μM 5,6-Dichloro-1-β-D-ribofuranosylbenzimidazole (DRB) for 2h) or with the corresponding volume of DMSO.

For BioID in HEK293T cells, pWPIeGFP (negative control), BioID:LCKN18 (positive control), BioID:TPR and pLgwBirAmChDamNup153 were transfected using JetPRIME (Polyplus) as instructed by the manufacturer's protocol. For BioID and DamID, small-scale production of lentiviruses was generated in HEK293T cells using JetPEI (Polyplus) and psPAX2 (a gift from Didier Trono), pVSV-G (a gift from Akitsu Hotta), pAdVAntageTM (Promega) and specific vectors in the ratio of 1.1 μg, 2 μg, 0.3 μg and 3.1 μg as instructed by the manufacturer's protocol. Supernatant containing virus particles was collected after 48h-72h and 1 ml was added onto 5e6 of Jurkat cells before spinoculation at 2300 rpm for 90 min.

For BioID, 50 μM biotin (Sigma-Aldrich) was added 24 h post transfection/transduction. DamID and BioID transduced cells were collected 72h post transduction.

For shRNA experiments, lentiviruses were produced in greater scale in HEK293T cells using psPAX2, pVSV-G, pAdVAntageTM (Promega) and pLKO.1 cloning vector (a gift from David Root) in the ratio of 58.4 μg, 32 μg, 9.2 μg and 90 μg.

Plasmids were prepared in 4 ml of OptiMEM (Gibco) and mixed with 4 ml of OptiMEM containing 400 μl of PEI transfection reagent, incubated at room temperature for 30 min, and 2 ml were pipetted onto four 15cm-dishes (5e6 cells/dish). The supernatant containing virus particles was collected after 48h-72h and filtered via 0.45 μm filter (Roth/Millipore). Virus was concentrated using 20% sucrose and ultracentrifugation at 24,000 rpm (Beckman SW28 rotor) for 2h at 4 °C. The supernatant was discarded and virus was resuspended in 200 µl fresh PBS with 0.1% FCS for 30 min at 4°C, aliquoted and stored at -80 °C. Virus titers were assessed by determination of reverse transcriptase activity (SG-PERT). 5 x $10^6$ Jurkat cells were infected with 30 ng lentivirus using spinoculation at 37 °C with 2300 g. Three days post infection, cells were selected with 1 µg/ml puromycin. Cells were then assessed for viability (MTT, Promega), pelleted for RNA extraction, western blot or fixed onto coverslips (IF/PLA) or µ-slide 8-well glass bottom chambers (FISH, IBIDI) .

**RNA extraction, RT-qPCR and PCR**

RNA was extracted from pelleted cells with Nucleospin RNA (Machinery-Nagel) according to manufacturer instructions. 0.5 µg of RNA quantified by nanodrop was then retrotranscribed using MultiScribe Reverse Transcriptase kit (Applied Biosystems) with random primers, dNTPs and RNase inhibitor (Roche or Thermo Fisher) added to the retrotranscription buffer provided by the kit. Protocols have been followed according to manufacturer recommendations. For the Real Time qPCR, 2 µL of cDNA were used upon dilution (1:10) and the amplification was performed with SsoFast EvaGreen Supermix (Biorad). Primers depicting exon/exon or intron/exon amplicons are reported in Table 1. Intron retentions was further assessed using PCR with exon primers, designed to amplify a region across two exons. cDNA from whole cell RNA was amplified with exon primers (Table 1) using 2X MyTaq Master Mix (Meridian Bioscience™). Afterwards, PCR products were visualized on a 2% agarose gel to detect different sized cDNA fragments.

**Co-Immunoprecipitation (Co-IP)**

Cells were washed once and scraped in PBS supplemented with cOmplete™ Mini EDTA-free protease inhibitor cocktail tablets (PBS+). Cells were pelleted and lysed for 15 minutes spinning incubation at 4 °C in RIPA buffer (20 mM Tris HCl pH7.4, 150 mM NaCl,1 mM EDTA, 0.5% NP40, 0.5% Sodium deoxycholate, 0.1% SDS + cOmplete EDTA-free miniProtease Inhibitor cocktail tablet (Roche) followed by sonication on a Bioruptor™ Plus Sonicator (12-15 cycles, 30 sec on/30 sec off). Protein content was measured with micro BCA protein assay kit (Pierce) and 500 µg of lysate was incubated overnight with 25 µL ChromoTek GFP-Trap® Agarose beads slur (Proteintech Lab). Co-immunoprecipitation was performed as per manufacturers protocols. Immunoprecipitated material was eluted in 50 µL of Laemmli buffer, boiled at 65°C for 30 min and 95°C for 5 min before SDS-PAGE.

**SDS-PAGE and Western blot**

Cells were rinsed twice in PBS and lysed with RIPA buffer before sheared in a water bath sonicator 3-5 cycles or until viscosity disappearance. Quantification of protein concentration was performed using micro BCA protein assay kit (Pierce) and equal amount of protein conditioned with lane marker reducing sample buffer 5X (Pierce), incubated 5 min at 95 °C and loaded in a precast NuPAGE Bis-

Tris 4-12% SDS-PAGE. Protein lysate was transferred to a nitrocellulose membrane with a wet-transfer system (Biorad), overnight at 30V 4 °C.

Ponceau staining was performed to visualise the efficiency of protein transfer to the membrane that was then blocked for 1 hour at room temperature with 5% milk in PBST. The incubation with primary antibody was performed either at 4 °C O/N or at 1 hour at room temperature. Primary antibodies used available in **Supplementary Table 2**. After washes, the membrane was probed 1 hour room temperature with secondary antibody conjugated with horseradish peroxidase (HRP). Once washed and incubated with ECL substrate (Pierce), the membrane was analyzed with ECL chemostat (iNTAS).

**BioID**

For each experiment, 20e6 snap-frozen cells were used and treated as PMCID: PMC5740495. In short, pellets were resuspended in 5mL lysis buffer (50 mM Tris pH 7.5, 150 mM NaCl, 1 mM EDTA, 1 mM EGTA, 1% Triton, 1 mg/ml aprotinin, 0.5 mg/ml leupeptin, 250U turbonuclease (Accelagen), 1 mM PMSF and 0.1% SDS) for 1 hour in a rotator before sonication until viscosity was absent. Lysate was centrifuged before adding 45 µL of streptavidin beads slur (GE life sciences) for 3h at 4 °C. The beads were transferred to a spin column, and the digested peptides were eluted twice with 50 mM ammonium bicarbonate. To remove the biotinylated peptides still bound to the beads, 150 µl of 80% ACN and 20% TFA was added, briefly mixed, and eluted; this step was done twice. The ACN/TFA elutions were merged. The samples were dried using a vacuum centrifugation. The elutions were resuspended in 200 µl buffer 5% ACN, 0.1% Formic Acid. The desalting and clean-up of the samples were carried out using Micro Spin Columns (Harvard Apparatus). The shot-gun MS experiments were performed as previously described PMCID: PMC5740495.

**BioID Data analysis**

In total 5 datasets were analysed. Triplicates were generated in HEK293T included BirA fused to Nup153, TPR and LCKN18 (positive control, provided by the laboratory of Oliver T Fackler) and untagged eGFP (negative control). Duplicates were generated in Jurkat T cells including BirA fused to TPR or untagged eGFP (negative control). Filtering of BirA fused proteins was based on the number of peptides (>2), LFQ (>5th quantiles of all values) and presence of corresponding peptides and LFQ in the GFP negative control (<3rd quantiles of all negative control values). Three protein lists were created. The first list encompassed the peptides found in the 3/3 HEK293T datasets whilst the second encompassed the peptides present in 2/3 of the HEK293T datasets. Finally, the third list recovers peptides present in only 1/3 of the HEK293T datasets but that match at least one valid value in the Jurkat data set. This extra filtering step allows for a controlled loss of information while still extracting possibly interesting information from the HEK293T data set. After combining all three lists, repeated entries were removed, generating a final list with 638 entries. DAVID and STRING were used for Gene ontology and network analysis, respectively. The STRING network was created using Kmeans with 6 clusters and high confidence interaction score.

**Immunofluorescence, proximity ligation assay and SABER FISH**

Cells were grown in poly-lysine coated coverslips (HEK293T) or seeded onto coverslips/µ-slide 8-well glass bottom chambers pre-treated with 1 mg/mL PEI (Jurkats). Cells were fixed with 4 % Paraformaldehyde (PFA) in PBS for 10 min at room temperature; 0.5 % Triton X-100/0.1% Tween-20 in PBS was added in the final 2 min. In case of Jurkat cells, cells underwent hypotonic shock for 30 sec in 0.3x PBS before fixation. After fixation cells were washed three times in PBS and permeabilized in 0.5 % Triton X-100 in PBS for 10 min. After another three washes in PBS coverslips were used directly for further experiments or stored in 20 % glycerol in PBS at 4 °C. For SABERFISH, cells were processed immediately after fixation.

**Immunofluorescence**

For immunofluorescence, coverslips with attached cells were removed from glycerol, transferred to a humidity chamber, and washed in PBS. Then, coverslips were blocked in 4 % bovine serum albumin (BSA) for 30 min at room temperature and were incubated in primary antibodies (in 4 % BSA) overnight at 4 °C (**Supplementary Table 2).** After incubation, coverslips were washed three times in PBST 5 min each. Coverslips were incubated in AlexaFluor-or ATTO-conjugated secondary antibodies in 4 % BSA for one hour at room temperature. After another three washes (5 min in PBS), nuclei were visualised with Hoechst 33342 (1:10,000 in PBS). Finally, coverslips were dipped into water, dried on filter paper, and mounted on a glass slide with Mowiol (Merck Millipore).

**In-situ PLA**

The *in-situ* PLA is a biochemical assay designed to detect the interaction of two proteins, which are in a proximity of at least 40 nm (*3*). We performed the assay as instructed by the manufacturer's protocol using the anti-mouse MINUS and anti-rabbit PLUS Duolink® In situ PLA® probes with the Duolink® in-situ detection reagent red. Before starting *in-situ* PLA, working dilutions for used antibodies were confirmed in IF staining. Briefly, coverslips were blocked in 1x Duolink® blocking solution for one hour at 37 °C in a humidified chamber. Blocking solution was removed and primary antibodies (see below), diluted in 1x Duolink® antibody diluent, were pipetted on the coverslips.

After overnight incubation at 4 °C, coverslips were washed three times 5 min each in Duolink® *in-situ* wash buffer A and then incubated in oligonucleotide-labeled secondary antibodies. Here, anti-mouse MINUS and anti-rabbit PLUS Duolink® In situ PLA® probes were mixed 1:5 in 1X Duolink® antibody diluent and incubated on the coverslips for one hour at 37 °C in a humidity chamber. After three washes, oligonucleotides were ligated with a ligase (0.025 U/µl) in 1x ligation buffer in water for 30 min at 37 °C. Coverslips were washed again followed by rolling circle amplification, initiated with

0.125 U/µl polymerase in 1x amplification red buffer. After final washes in Duolink® *in-situ* wash buffer B (3x 10 min in 1x wash buffer B and single wash in 0.01x wash buffer B in water), coverslips were mounted on glass slides in Duolink® *in-situ* mounting medium with DAPI. Coverslips were sealed using nail polish and imaged the following day. To assess Nup153 levels in Nup153 KD conditions, *in-situ* PLA and immunofluorescence staining were combined. After washing in wash buffer B, blocking and incubation in primary (Nup153; ab24700) and secondary antibody was performed as described above. After final washing steps (3 x PBST for 5 min, 1 x wash buffer B for 10 min and a single wash in 0.01x wash buffer B), coverslips were mounter on glass slides in Duolink® *in-situ* mounting medium with DAPI. Coverslips were sealed using nail polish and imaged the following day.

**SABER FISH**

For SABERFISH (*4*), IBIDI chambers were washed with 2xSSC supplemented with 0.1% Tween prior to blocking with 1 µM random 20mer in hybridization mix (10% Dextran sulfate, 0.1% Tween-20, 50%Formamide in 2xSSC) for 3 min 60 °C followed by 7 min at 40 °C. 25nM concatemer pool were hybridized in hybridization mix 4 min 60 °C followed by overnight at 42 °C. IBIDI chambers were washed four times with 2xSSCT, pre-warmed to 60 °C, and two times at room temperature. IBIDI chambers were then blocked with 4% BSA for 30 min and stained for Nup153 (ab24700) overnight at 4 °C or for 2h at room temperature. After washing with PBST, Alexa secondary mouse 568 was added for 1h (1:1000) and chambers washed with PBST before imager hybridization. For that, 10nM imager was added in PBS for 1h and washed in PBS before DAPI/Hoechst 33342 (1:10,000 in PBS) staining. Chambers were washed in PBS and imaged.

Probes were designed using OligoMiner, as previously described (*5*)

**Microscopy**

Images were taken on Leica TCS SP8 DLS microscope, Zeiss LSM900 confocal microscope equipped with Airyscan Detector or Expert Line STED system with an SLM based easy3D module.

Fluorescence from IF, *in-situ* PLA and SABERFISH was measured in a z-stack using Zeiss LSM900 confocal microscope equipped with Airyscan Detector and a Plan-Apochromate 63x/1,4 oil immersion objective. Furthermore, IF staining was evaluated with super resolution mode using Airyscan detector.

STED imaging was performed on an Expert Line STED system with an SLM based easy3D module using a 100x oil immersion objective (NA. 1.4). STED images were restored with

Huygens Deconvolution by using Classic Maximum Likelihood Estimation (CMLE) algorithm and Deconvolution Express mode with "Conservative" or "Standard" setting.

**Image Analysis**

To segment PLA signals we trained a Random Forest classifier using ilastik (Berg et al., 2019) pixel classification workflow. The training set of data was arbitrarily selected and very sparsely labelled (<0.1% of total pixels were manually categorized into "signal" and "background" categories). For nuclei segmentation (Hoechst staining), a model based on deep learning algorithms was trained employing the arivis Cloud platform (www.apeer.com). The same models were used for signal segmentation across all experiments. This approach ensured an unbiased PLA and nuclei signal segmentation across all experiments. The distance of PLA/SABERFISH signals from the nuclear envelope was measured in Imaris. Therefore, segmented channels were merged, and the resulting z-stacks were introduced into Imaris. The nuclear signal was transformed into a surface object (0.2 smooth factor) from absolute intensities. PLA/SABERFISH spots were processed into spot objects with XY diameter 0.5 µm and 1 µm Z model PSF elongation. The object-object statistics allow the calculation of the nuclear volume and the closest distance between PLA spots and the nuclear surface. All the measurements were done in the 3D space. Only PLA/SABERFISH signals whose centroid was located within the nucleus were considered and the number of signals per nucleus was determined. The distances were normalized by the mean nuclear radius to derive relative distance to the closest nuclear edge which were binned and plotted in a histogram (bin size: 0.1) in GraphPad Prism9.

STED images were analyzed in Imaris. The DAPI channel was used to create a surface with 0.2 smooth factor. Distribution of the splicing proteins and the Nups were assessed by applying the spot function (XY diameter 0.1 µm and 0.2 µm Z model PSF elongation) on the respective channel. After filtering out spots that were assigned to low intensities, spots inside the nuclear surface were selected and the closest distance to the surface was calculated. For splicing proteins, also the closest distance to Nups spots was calculated. Again, relative distances were created by normalization to nuclei radius and frequency distribution was plotted in a histogram (bin size: 0.1).

**DamID-Seq**

DamID was performed as described in (Vogel et al., 2007). Briefly, triplicate experiments were prepared from 5 million Jurkat cells transduced with 1mL of filtered lentivirus produced with either

DAm-only or Dam fused to Nup153 using spinoculation. After 72h cells were pelleted and subjected to DNA extraction, using the Bioline Isolate II genomic DNA kit including proteinase K and RNAseA digestion as per manufacturer's instructions. 0.5 µg of extracted DNA was then digested by DpnI (NEB) and, following heat inactivation of DpnI, was ligated to the DamID adaptor duplex (dsAdR) generated from the oligonucleotides AdRt (5'-CTAATACGACTCACATAGGGCAGCGTGGTCGCGGCCGA-GGA-3') and AdRb (5'-TCCTCGGCCG-3') after which DNA was further digested by DpnII (NEB). To amplify DNA sequences methylated by the Dam methylase, 8 µl of DpnII digested material was then subjected to PCR using the Bioline MyTaq polymerase and in the the presence of the 1.25 µM Adr-PCR primer (5'-GGTCGCGGCCGAGGATC-3') modified to contain random nucleotides in the beginning of the sequences ("N") providing sample heterogeneity. Each sample Dam-only/DamID:Nup153 (in triplicates) was amplified with:
NGGTCGCGGCCGAGGATC,
NNGGTCGCGGCCGAGGATC,
NNNGGTCGCGGCCGAGGATC,
NNNNGGTCGCGGCCGAGGATC or
NNNNNGGTCGCGGCCGAGGATC.
Following PCR, the distribution of amplified DNA fragments was checked on agarose gels. 500ng of PCR product was purified using Ampure Beads before library preparation using NEBNExt Ultra II kit (NEB) and profiled by Bioanalyser (Agilent) and then mixed to achieve equimolar amounts of each library. DNA was sequenced on a NEXTSeq550 to produce single-end 150bp reads, according to manufacturer instructions.

**Statistical Analysis**

Details of statistical analyses, including the number of individuals and nuclei in imaging experiments, as well as the statistical methods employed, and P values, are provided in each figure legend, the text, or Supplementary Tables. The results presented in this article are based on data obtained from a minimum of three individuals, with exceptions specified in figure legends. All statistical analyses were conducted using Prism software (GraphPad Software; SCR_002798). Significance was determined using paired or unpaired t-tests (one or two-tailed), ordinary Two-way analysis of variance (ANOVA), and ordinary One-way ANOVA. Results were considered significant when the P value was <0.05.

**Bioinformatic analysis**

**DamID analysis**

Each individual replicate contained between 15-25M PE reads and were processed as in (*1*). Briefly, data quality was assessed with FastQC version 0.11.9

(http://www.bioinformatics.babraham.ac.uk/projects/fastqc/), adaptors removed with Trimmomatic (version 0.35) and aligned to the human genome build Hg19 with BWA MEM (version 0.7.16) https://github.com/lh3/bwa
using default parameters. Mitochondrial sequences (< 1%) and non-primary alignments were filtered out, and duplicates marked with samtools markdup (version 1.17)(*6*) . To identify Nup153 peaks, replicates were compared for reproducibility and then pooled. Quantitation was performed at the DpnI fragment level separately for the control 'dam-alone' and the Nup153-dam samples, and then normalised to the library size. Next the log2(Nup153-dam/dam) was calculated, after adding a pseudocount of 0.001 to every bin in order to avoid division by zero errors. The resulting bedGraph was converted to bigwig format using wigToBigwig version 377 for visualisation on a genome browser https://www.encodeproject.org/software/wigtobigwig/. Peaks were obtained with MACS2 (version 2.2.7.1) in broadPeak mode with parameters –nomodel –extsize 360 –shift 180, and further filtered by FDR 0.01%, fold-enrichment > 3-fold, and minimum peak length of 1Kb. In addition, peaks were screened for their proximity to a LAD using the Jurkat LAD dataset (*1*), marking as LAD-proximal any peak that is < 100 Kb from a LAD and at least 10 Kb away from a genomic sequencing gap. The genome gaps file for Hg19 was retrieved from the UCSC's table browser tool (https://genome.ucsc.edu/cgi-bin/hgTables).

**Annotation of Nup153 peaks on the genome**

For annotation of Nup153 peaks on the genome, the Nup153 locations/peaks were loaded into R in .bed file format and were annotated using UCSC annotation databases using the TxDb.Hsapiens.UCSC.hg19.knownGene package (version 3.2.2.). Pie charts were created using the chipSeeker package (version 1.38.0)(*7*).

**Intron length and GC-content of Nup153 bound genes**

10 fully random samples of genes (n=500) were chosen from genes available in Ensembl databases using the biomart R package (version 2.58.2), in order to compare their intron length and GC content to Nup153 associated genes. Biomart was then used to get the data for all transcripts and all exons for each chosen gene, since Ensembl doesen't store intron information directly.

The data was filtered to only keep the longest transcript for each gene (the one with the greatest number of exons, and the intron start and stop locations were calculated by assigning the first intron start to be the first exon end, and the first intron end to be the start of the second exon etc.

For the GC content of the introns the packages BSgenome.Hsapiens.UCSC.hg19 (version 1.4.3) and Biostrings (2.70.2) were used to get the DNA composition of the above calculated intron coordinates from the ful genome sequence hg19 stored by UCSC.

**LADs definition and analysis**

Discrete LAD and inter-LAD regions were identified from resting Jurkat cells using Lamin B1-DamID data (Robson et al., 2017) and a two-state Hidden Markov Model (HMM) (https://github.com/gui11aume/HMMt). For Figures 4f and 4g, the genomic distance between the center of each Nup153 peak and the nearest LAD border was calculated using custom Python code. As a control, 5 bootstrap samples of 10,000 genomic coordinates (peaks) were randomly generated, maintaining the same LAD/inter-LAD ratio as the Nup153 peaks obtained by DamID.

**Nup153 data analysis GPSeq**

Genome-wide radial positioning was estimated using GPSeq scores from HAP1 cells (*8*). To assign a GPSeq score to Nup153 peaks, each peak was intersected with genomic windows of 1 Mb with a 100 kb step size, and the GPSeq scores were averaged. For visualization, GPSeq scores were transformed using 1-log2(GPSeq) to denote 1 as the periphery. Figures were generated using custom Python and matplotlib code.

**Chromatin data - sources:**

**H3K4me3** - GSM569085; **H3K27ac** - GSM1697882; **H3K36me3** - GSM1603209; **H3K27me3** - GSM1062740; **RNAPII S5P** - GSM1603223; **RNAPII S2P**-GSM1603221; **Brd4** - GSM1961565; **H3K4me1** - GSM1220567; **H3K79me2** - GSM569089; **H3K9me2** (*9*)

**Chromatin data - analysis:**

Raw ChIP-Seq data was downloaded from SRA. Reads were trimmed, mapped and deduplicated using programs from BBTools package, version 38.94 (BBduk2, BBMap, and Dedupe respectively) (https://sourceforge.net/projects/bbmap/). Resulting bam files were normalized with RPKM method using deepTools' bamCoverage (Ramírez et al., 2016)

**LADs analysis with respect to chromatin marks:**

To define LAD regions with respect to chromatin marks, we started with LADs from GSE9497 (*1*). Constrained release of lamina-associated enhancers and genes from the nuclear envelope during T-cell activation facilitates their association in chromosome compartments(*1*). Since these regions appeared to be highly fragmented, we decided to re-define LADs for the purposes of this work in order to get better LAD definition with respect to chromatin marks. We took into account additional data: LADs from Guelen et al (*11*), and chromatin marks Brd4 (GSM1961565), H3K27ac (GSM1697882), H3K36me3 (GSM1603209), H3K4me1 (GSM1220567), H3K4me3 (GSM569085), H3K79me2(GSM569089), H3K9me2 (*9*), H3K27me3 (GSM1062740). We choose those chromatin modifications because Brd4, H3K27ac, H3K36me3, H3K4me1, H3K4me3 and H3K79me2 are expected to be negatively correlated with LADs, whereas H3K9me2 and H3K27me3 should be positively correlated (Guelen et al., 2008).

For each of chromatin marks, we calculated average coverage in 10 kb tiles from RPKM-normalised files. We marked LAD-covered areas with 1 and areas without LADs with 0, and again calculated the average in 10 kb tiles. We assigned 2 points to areas with LAD signal for each of LAD datasets, and 1 point for each "correct" chromatin signal (positive coverage for H3K9me2 and H3K27me3, and negative for the rest, with weakest 5% of signal excluded from analysis). Tiles which got assigned 7 points or more were merged together if distance between them was 100 kb or less. Resulting LAD areas shorter than 10 kb were excluded, as well as areas positioned in largest 25% of blacklisted regions according to ENCODE blacklist (*12*)

**Plotting**

Enrichment plots were generated using ngsplot, version 2.63 (*13*) with default parameters. Plotted lines represent average signal across all Nup153-bound genes, all genes intersecting with LADs, and all other protein-coding genes which don't fit into first two categories.

**References related to Supplementary Figuger, Figure legends and Materials and Methods:**
